# Enhancing Medical Science Engagement Among Medical Undergraduates Through International Research Exchange

**DOI:** 10.1101/2025.02.26.640317

**Authors:** Monika Jürgenson, Andrea García-Llorca, Anu Sarv, Thor Eysteinsson, Miriam A. Hickey

## Abstract

**Background:** The global decline in the number of physician-scientists, despite an increase in practicing physicians, underscores a critical need for integrating research training into medical education. Addressing this issue, we established an international research exchange program aimed to enhance scientific literacy, foster transferable skills, and align curricula with European standards through collaborative research experiences.

**Methodology:** The program enabled reciprocal student mobility, involving eleven medical undergraduates who conducted month-long basic science research projects. Participants also completed comprehensive pre-training in scientific communication, safety protocols, and ethics, and were required to participate in local public engagement events. Feedback from participants, which we present here, was collected via three anonymous, voluntary questionnaires: pre-program, post-program and post-1-year follow-up, which we provide here to support similar initiatives.

**Results:** Despite challenges and delays due to the COVID-19 pandemic, the program met its objectives, demonstrating adaptability and effective resource management. Feedback revealed significant improvements in participants’ confidence in research methodologies, critical appraisal of scientific literature, and motivation for future research involvement.

**Conclusion:** This project highlights the potential of structured international exchange programs, particularly among smaller institutions, to address gaps in medical education, enhance scientific training and opportunities in translational research for undergraduates, and cultivate the next generation of physician-scientists.

## 1 Introduction

The number of physician-scientists is declining globally, despite the overall rise in the number of practicing physicians (1). This concerning trend has prompted national and international medical organizations to advocate for increased engagement in research. For example, national guidelines in Canada and the United States encourage physician participation in research, while the 2024 draft of the UK’s Good Medical Practice explicitly highlights research as a core skill within medical training(2– 4). Critical thinking and evidence-based approaches have long been recognized as fundamental skills for healthcare professionals, further underscoring the need to address the decline in science-based skills (5,6).

While studies have shown the benefits of international exchanges for medical students, many primarily focus on the development of clinical skills (7–15). With respect to science skills, barriers to participation exist, particularly for undergraduate medical students. Time constraints, a lack of appropriate research projects, and the exclusion of research from standard curricula remain significant obstacles(16). Additionally, smaller countries face further challenges due to limited laboratory resources, expertise, and opportunities(17), hindering their ability to attract and nurture young talent in both medicine and research (17,18).

Participating in real-world, structured translational science projects has been shown to increase the likelihood of medical students engaging in subsequent research activities (19–21). Moreover, international exchanges offer broad benefits for medical students, although most existing programs focus on clinical skill development rather than basic science research. Programs like the International Federation of Medical Students Associations’ (IFMSA) SCORE initiative highlight the potential of laboratory-based research exchanges to complement medical education by fostering critical thinking, technical skills, and global awareness.

Recognizing these gaps and opportunities, we established a research exchange program between two partner universities aimed at undergraduate medical students. By facilitating month-long, hands-on research projects, this initiative provided students with practical exposure to basic science, enhanced transferable skills, and enriched the academic curricula of both institutions. The program also aligned with the Bologna Process goals, fostering international collaboration and strengthening the competitiveness of the participating universities.

Through careful planning and execution, the program addressed common barriers to student participation in research. It included pre-training in communication, safety, and ethics, and required students to complete all laboratory hours and submit detailed reports to earn academic credit. This model not only ensured rigorous academic standards but also created a framework for mutual learning and cultural exchange.

The outcomes of this program underscore the potential of structured international research exchanges to enhance medical education. By fostering a deeper understanding of science and its applications, such initiatives can play a crucial role in reversing the decline of physician-scientists and preparing the next generation of healthcare professionals to meet the challenges of modern medicine. Based on our experience, we also propose practical guidelines for establishing similar programs to broaden their impact and reach.

## 2 Materials and Methods

### 2.1 Participants and Ethical Considerations

Medical undergraduate students were invited to participate in an international exchange program to conduct month-long basic science research projects (“research miniprojects”). Participation in feedback was voluntary, and students provided anonymous feedback through three questionnaires (Supplementary materials 1, 2 and 3) administered at different stages of the program. The questionnaire content was developed based on requirements identified by the authors (Monika Jürgenson, Andrea García-Llorca, Anu Sarv, Thor Eysteinsson and Miriam Hickey) and guided by relevant published commentary (22). The program was conducted in 2019-2023. For experiments involving animals, all necessary ethical approvals were obtained from local authorities before the students’ arrival. Ethics approval for the program surveys was granted by the Research Ethics Committee of the University of Tartu (approval number 361/T-8).

### 2.2 Program Design and Online Learning Platform

A dedicated online learning platform was created within Moodle space to support the participants. The platform provided general program information; introductory details about faculty members; mandatory training modules on laboratory safety, research animal ethics, and communication skills, background information on research projects, including updates reflecting new data as the projects evolved, templates and guidelines for progress and final reports, and finally resources on open science practices and our end-of-program conference. All students completed mandatory quizzes on safety and ethics and provided signed Laboratory Safety Guidelines before travel. Training on research animal ethics was mandatory for all participants, along with two academic hours of communication training focused on addressing diverse audiences, including peers, senior faculty, and the public. Finally, all students were required to participate in local public education events.

### 2.3 Research Miniprojects

Each student participated in an individual research miniproject as part of the exchange program. During these exchanges, students from the coordinating institution conducted research in the partner institution, while students from the partner institution carried out their projects in the coordinating institution. Students explored diverse research topics, including retinal morphology and degeneration, obesity, Alzheimer’s disease and cognitive performance, Wolfram syndrome, Parkinson’s disease, retinal function, and lysosomal function. This enabled our students to contribute to advancements in very different biomedical fields that were highly translational in nature.

### 2.4 Data Collection and Feedback Mechanism

Feedback from participants was collected via three questionnaires: a pre-program questionnaire: administered before the research exchanges to assess expectations and preparedness; a post-program questionnaire: administered immediately after program completion to evaluate satisfaction, challenges, and outcomes; and post-1-year follow-up questionnaire: designed to assess the long-term impact of the program on participants’ academic and professional development.

Exact questionnaires are presented in supplementary materials (Supplementary materials 1, 2 and 3). Answers to questions were based upon a five-point Likert scale, or free text.

### 2.5 Statistical analyses

Statistical analyses and graphical presentations were performed using GraphPad 9.4.1 (San Diego, CA, USA). All responses from student participants were considered when interpreting the results. In the pre-program questionnaire, 10 out of 11 students responded. The post-program questionnaire received responses from 9 out of 11 students, while the one-year follow-up questionnaire had 4 responses out of 11. Data sets were analyzed using Fisher’s exact test, with statistical significance set at p < 0.05.

We used G*Power to determine sample sizes. Based upon an initial even spread of students within each category, thus 20% of students classifying themselves within poor, fair, average, good or excellent answers to Likert-type questions, and that following the research miniproject, students’ perceptions changed to good (50% of students) or excellent (50% of students), N=8 students would be required in total (effect size 0.9, alpha = 0.05, power = 0.8, Df = 4). In a more conservative assessment, based upon an initial even spread of students within each category, thus 20% of students classifying themselves within poor, fair, average, good or excellent, and that following the research miniproject, students’ perceptions changed to average (20% of students), good (40% of students) or excellent (40% of students): N=15 students would be required in total (effect size 0.9, alpha = 0.05, power = 0.8, Df = 4).

## 3 Results

### 3.1 Laboratory skills

As presented in Figure 1A, feedback-based assessment demonstrated a statistically significant enhancement in technical science skills (p<0.0001, Fisher’s exact test). Critical assessment of scientific publications also improved but this was non-significant (Figure 1B). The findings suggest that practical laboratory training, lasting just one month, may improve ability to read basic science publications, and that it contributes positively to hard-skill development.

**Figure 1.**
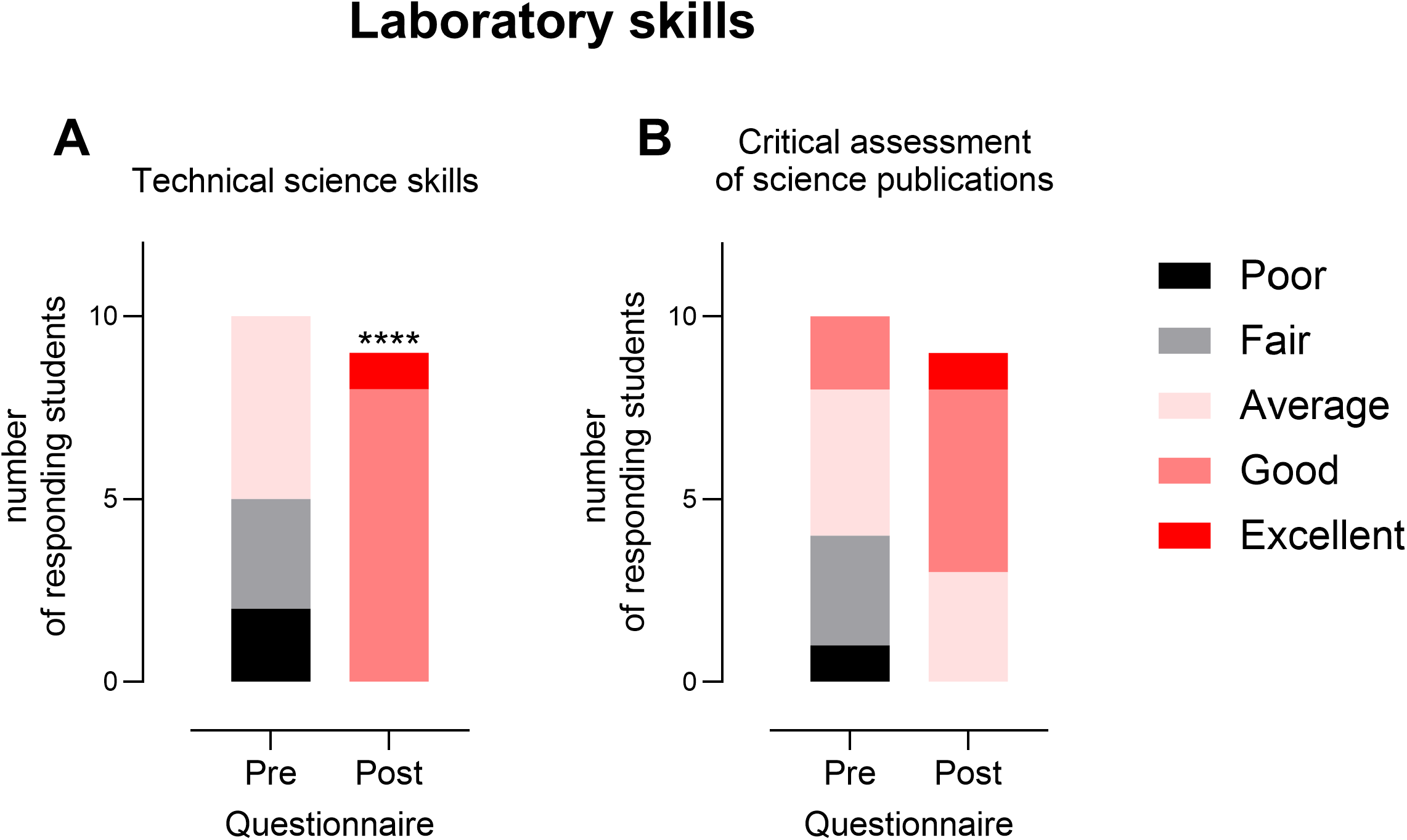
Laboratory skills The effect of the program on the laboratory skills of the participants. (A) Technical science skills. **** p < 0.0001; Fisher’s exact test; (B) Critical assessment of science publications. Number of participants Pre-questionnaire: n =10; Post-questionnaire: n = 9.

### 3.2 Transferable skills

As presented in Figure 2, students initially considered their transferable skills as good, for example, most students considered their time management, organisational skills, ability to work in teams and leadership skills as good. However, communication skills were rated as average-good and writing reports was considered to be average (Figure 2A-F). After participating in our program, we observed improvements across all of these outcomes, with students considering their skills as good-excellent for time management, organisation, working in teams and in communication and writing reports. Indeed, there was a significant improvement in communication skills (Figure 2E, p<0.05, Fisher’s exact test) and in writing reports (Figure 2F, p<0.01, Fisher’s exact test). Thus, just 1 month of experience in the laboratory improved several soft skills required of medical professionals(4,23).

**Figure 2.**
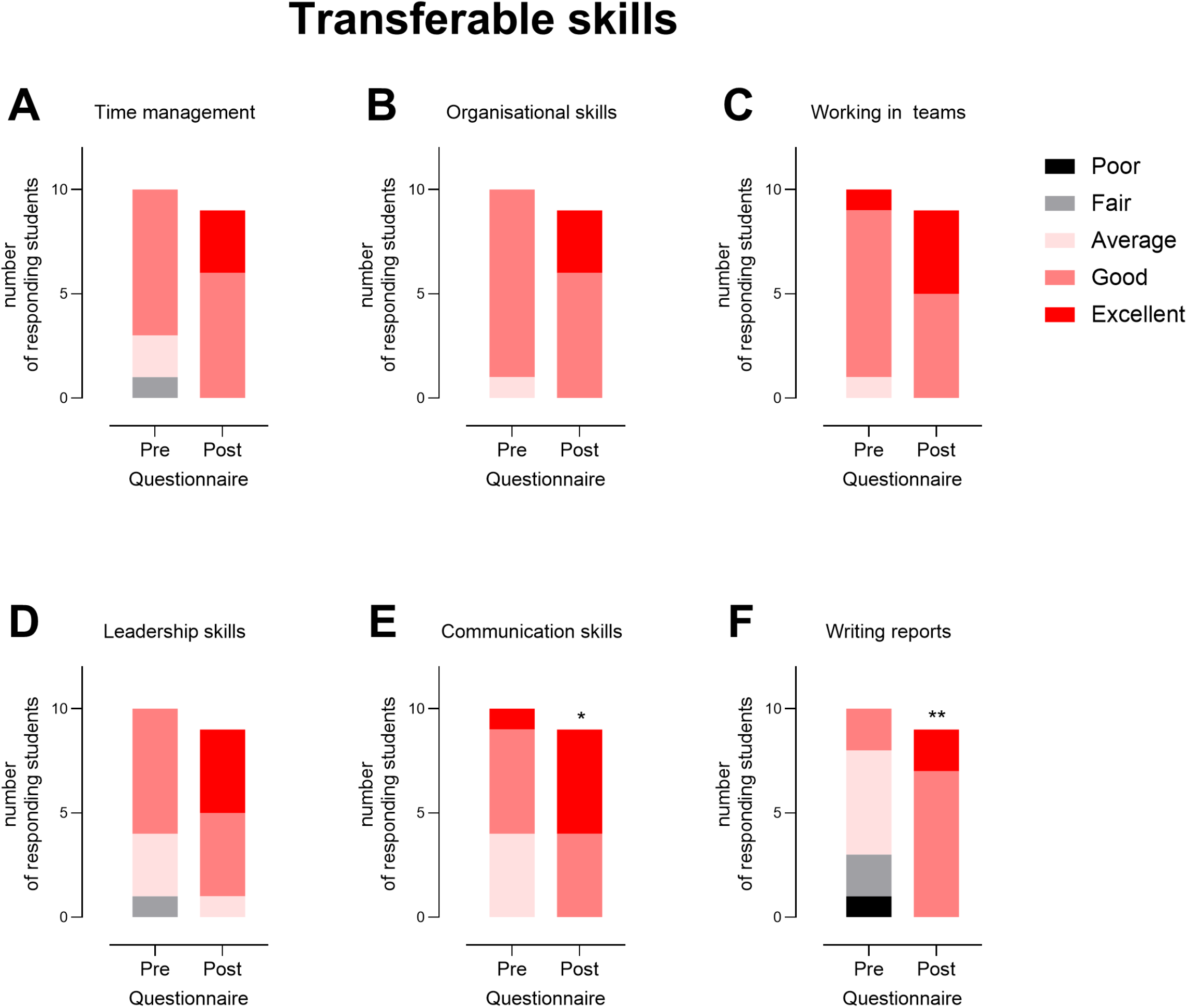
Transferable skills The effect of the program on transferable skills of the participants. (A) Time management skills; (B) Organizational skills; (C) Teamwork skills; (D) Leadership skills; (E) Communication skills. * p < 0.05; Fisher’s exact test; (F) Scientific report writing skills. ** p < 0.01; Fisher’s exact test. Number of participants Pre-questionnaire: n =10; Post-questionnaire: n = 9.

### 3.3 Overall opinion

We also asked participants to provide feedback on the overall impact of the program to better understand their perceived personal and professional development. Critically, as shown in Figure 3, students reported a large gain in understanding of how scientists tackle real-world problems (A). Students typically reported a large or very large gain in recognition of the importance of evidence in science (C). A moderate, large or very large gain was achieved in understanding ethical conduct in the field (D). Importantly, our program led to increased self-confidence among participants (E, large gain). There were more modest gains in their sense of contributing to the body of knowledge (B). This outcome is understandable, as undergraduate students may find it challenging to grasp all aspects of a foundational science project. Additionally, since the program involved a mini-project, completing it highlighted how much more there is to learn. Overall, participants reported feeling moderately to very satisfied with our program (Figure 3F).

**Figure 3.**
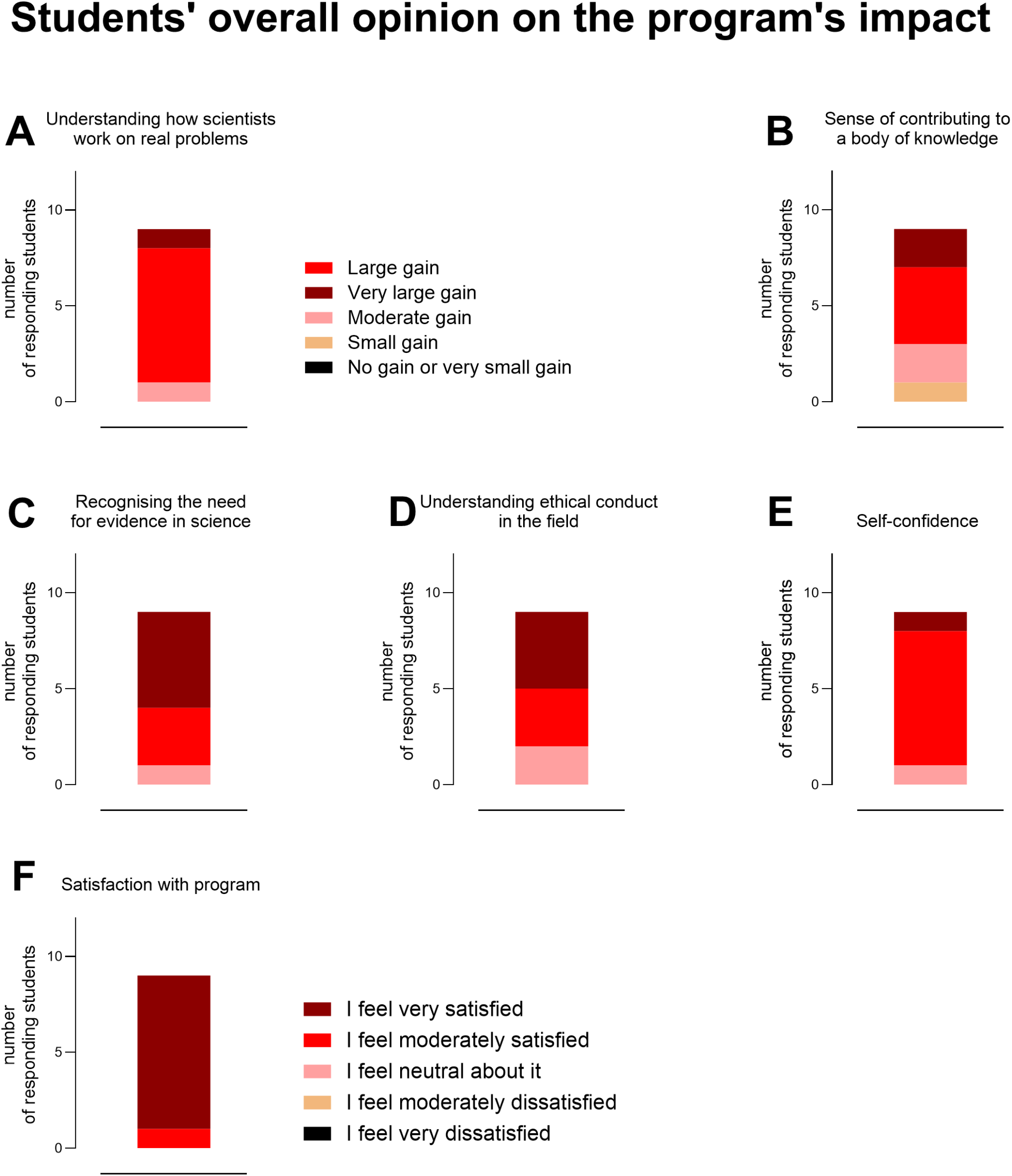
Students’ overall opinion on the impact of the program. Students’ overall opinion on the impact of the program. (A) Understanding of how scientists work on real problems, legend with A is for A - E; (B) Sense of contributing to a body of knowledge; (C) Recognizing the need for evidence in science; (D) Understanding ethical conduct in the field; (E) Self-confidence; (F) Satisfaction with the program. Post-questionnaire: n = 9.

### 3.4 Long-term impact of the program

A one-year follow-up questionnaire was conducted to evaluate the program’s long-term impact on participants’ academic and professional development. We received four responses: one participant is conducting a research project as part of their degree, while two are engaged in volunteer work-one of whom may transition into a degree-required project.

## 4 Discussion

Medical curricula are increasingly incorporating case-based learning to enhance relevance and real-world training and transitioning from basic science-based training (24). Most international electives available to medical students focus on clinical skills, while opportunities to develop laboratory and basic science skills remain limited. This is despite national organizations actively supporting and encouraging research training initiatives(2–4). Globally, physician-scientists face unique challenges in securing funding and training, leading to high attrition rates in sustaining a career that integrates both medicine and research (17). To mitigate this issue, some countries have introduced paid research training programs for medical graduates(17).

Surveys among undergraduate medical students indicate that barriers to engaging in research include time constraints, limited availability of suitable translational projects, and a lack of integration into the curriculum (19). However, those who participate report positive experiences (25). For smaller universities, international research exchanges offer significant benefits by expanding access to research opportunities and techniques. These programs also provide valuable transferable skills, such as proficiency in a common foreign language (English, in our case), which aligns with EU Recommendation on Key Competences for Lifelong Learning (2018/C 189/01). Additionally, such exchanges contribute to curriculum enhancement and support the Bologna Process by fostering student mobility and transnational collaboration (EU Bologna Process).

Our findings demonstrated significant improvements in laboratory and science skills among participants. Beyond technical skills, the program had a considerable impact on students’ transferable skills, as the post-program assessment revealed improvements in time management, organization, teamwork, and leadership, with statistically significant gains in communication skills and scientific report writing. These findings suggest that a hands-on research experience fosters essential skills beyond the laboratory, preparing students for interdisciplinary collaboration and effective scientific communication. A one-year follow-up questionnaire was conducted to evaluate the program’s long-term impact on academic and professional development and although out of the 11 eligible respondents, four provided feedback, they were all involved in science, whereas one participant was actively engaged in a research project as part of their degree. These results suggest that the program had a lasting effect on participants’ academic and professional trajectories, reinforcing the value of early exposure to research opportunities.

Our program showed the feasibility of conducting a research exchange program between two small countries on the borders of the EU, to the benefit of both. Importantly, institutional participants shared equal burden(7). Moreover, our program contributes to skills development required in recent guidelines placed by national bodies in the UK (4) and Canada (2) and also recommendations on healthcare electives (26–30).

While our findings highlight the benefits of international research exchange programs, some limitations must be acknowledged. First, the sample size was relatively small, which may limit the generalizability of our results. The number of participants decreased in the follow-up questionnaires, reducing the ability to assess long-term impacts comprehensively. Additionally, self-reported data may introduce response bias, as students might have felt inclined to provide positive feedback. Another limitation relates to variations in participants’ prior research experience. Differences in baseline skills may have influenced the degree of improvement observed, making it challenging to precisely quantify the program’s impact on individual skill development. Additionally, the short duration of the exchange (one month) may not have been sufficient for students to gain in-depth research expertise comparable to longer-term programs, although science-based skills were judged to have improved significantly. Logistical challenges also posed limitations. Although the program covered all expenses, accommodation costs varied between institutions, particularly since travel was most feasible during the summer months. Differences in accommodation costs, access to laboratory facilities in the host country, and funding disparities between partner institutions may have influenced the overall experience for some students. Additionally, external factors, including the COVID-19 pandemic and geopolitical instability, impacted program implementation and potentially influenced application rates. Despite these limitations, our study provides valuable insights into the potential of international research exchanges to enhance scientific training and skill development for medical students. Future programs could benefit from larger sample sizes and extended study periods. Indeed, here we provide guidelines based upon our experiences (Supplementary material 4) and building upon published recommendations (26–30) to support similar future programs between small countries.

## Conclusions

Overall, our findings highlight the significant benefits of international research exchange programs, offering hands-on research experience, strengthening both technical and transferable skills, and fostering global collaboration. Importantly, our results demonstrate how smaller EU border countries can collaborate to provide enhanced opportunities for students while equipping them with critical technical and transferable skills valued by international medical organizations. Many exchange programs are available within the EU. In particular, ERASMUS schemes are very well known and are very valuable for students and faculty alike. However, our program emphasized, specifically, collaboration between smaller countries, with a focus upon exposure to basic science. Addressing logistical challenges, ensuring structured training and supervision, and integrating participant feedback are essential for the continued success and expansion of such initiatives in the future.

## 5 Acknowledgments

We thank our student participants for their excellent work and input to our program. We thank Kadri Veeperv and Mari-Liis Timmotalo of the Medical Faculty of the University of Tartu and Magnús Guðmundsson of the Medical Faculty of the University of Iceland for their kind assistance with our program. We thank the student union of the University of Iceland for kind assistance with advertising our program.

## 8 Ethical approval and informed consent statement

Participants were provided with informed consent forms and invited to provide voluntary, anonymous feedback. Ethics approval for the program was granted by the Research Ethics Committee of the University of Tartu (approval number 361/T-8).

## 9 Data availability

The data underlying the results presented in the study are available in aggregated form and include the values used to build the figures. Individual participant responses, including anonymised data, are not publicly available due to ethical restrictions as approved by the Research Ethics Committee of the University of Tartu (approval no. 361/T-8). Requests regarding data access restrictions may be directed to the Research Ethics Committee of the University of Tartu.

This article was previously presented as a meeting abstract at the 2024 Annual University of Tartu Medical Faculty conference. This article was previously posted to the BioRxiv preprint server on February 28, 2025.

## 10 Author Contributions

Monika Jürgenson: Writing – review and editing, interpretation of results, conceptualization, supervision and funding acquisition Andrea García-Llorca: Writing – review and editing, interpretation of results, supervision. Anu Sarv: Preparation of questionnaires, interpretation of results, conceptualization. Thor Eysteinsson: Writing – review and editing, interpretation of results, conceptualization, supervision, funding acquisition. Miriam Hickey: Writing – review and editing, interpretation of results, conceptualization, supervision, resources, and funding acquisition.

