## Supplementary material 1 for "Enhancing Medical Science Engagement Among Medical Undergraduates Through International Research Exchange"

Based upon requirements identified by authors Monika Jürgenson, Andrea García-Llorca, Anu Sarv, Thor Eysteinsson and Miriam Hickey and upon published commentary (1).

#### Pre-program questionnaire

*Mode: Confidential*

| Question | Answer |
|---|---|
| How did you hear about this program? | (free text) |
| In one or two sentences, what do you understand by the term "scientist"? | (free text) |
| In one or two sentences, what do you understand by the term "physician-scientist"? | (free text) |
| In your opinion, what is the value of being a physician-scientist? | (free text) |
| Before you begin your miniproject, please rate your competency in the following areas. Information on the meaning of each area is provided underneath. | |
| Area | Rating |
| Technical science skills[a]<br><br>[a.] Your ability to perform an experiment in the laboratory. For example, during your miniproject, you will learn how to perform confocal microscopy or cognitive testing. | Poor |
| | Fair |
| | Average |
| | Good |
| | Excellent |
| Critical assessment of science publications[b]<br><br>[b.] Your ability to read a scientific paper and interpret and critique its data, for example a paper that provides background for your miniproject. | Poor |
| | Fair |
| | Average |
| | Good |
| | Excellent |
| Please rate your competency in the following areas. Information on the meaning of each area is provided underneath. | |
| Area | Rating |
| Time management[a]<br><br>[a.] Your ability to plan ahead to reach your targets, for example to complete a series of experiments in your miniproject. | Poor |
| | Fair |
| | Average |
| | Good |
| | Excellent |
| Organisational skills[b] | Poor |
| | Fair |

| | |
|---|---|
| [b.] Your ability to schedule your tasks and your punctuality, for example, to use a particular piece of equipment that you need for your miniproject. | Average<br>Good<br>Excellent |
| **Please rate your competency in the following areas.<br>Information on the meaning of each area is provided underneath.** | |
| Area | Rating |
| Communication skills[a]<br><br>[a.] Your ability to communicate effectively and efficiently with colleagues, your superiors and the public in written or oral format, for example, your ability to troubleshoot with your superiors and your ability to use technical and/or simple language to explain your data. | Poor<br>Fair<br>Average<br>Good<br>Excellent |
| Presentation skills[b]<br><br>[b.] Your ability to present data to your colleagues, your superiors, to the public. For example, at the end of the program, all students will present their data in the multinational meeting, and all students will present their data and their experiences to the public during European Science night. | Poor<br>Fair<br>Average<br>Good<br>Excellent |
| Writing reports[c]<br><br>[c.] Your ability to write a report that is accurate, fair and based upon evidence, for example, a report on the data that you obtain during your miniproject. | Poor<br>Fair<br>Average<br>Good<br>Excellent |
| **Please rate your competency in the following areas.<br>Information on the meaning of each area is provided underneath.** | |
| Area | Rating |
| Ability to work in teams[a]<br><br>[a.] Your ability to reach agreement on tasks to do within a team and ability to complete that task, for example, during the public presentation during European Science night. | Poor<br>Fair<br>Average<br>Good<br>Excellent |
| Leadership skills[b]<br><br>[b.] Your willingness to take on responsibility and organise team-based tasks, for example, during the public presentation during European Science night. | Poor<br>Fair<br>Average<br>Good<br>Excellent |

| | |
|---|---|
| You just rated your competency in several areas. | |
| Now please formulate your goals for your miniproject and this program, based on the areas that your feel you need to work on (refer to your ratings). | (free text) |

### Supplementary material 2

Based upon requirements identified by authors Monika Jürgenson, Andrea García-Llorca, Anu Sarv, Thor Eysteinsson and Miriam Hickey and upon published commentary (1).

**Post-program questionnaire**

*Mode: Confidential*

| Question | Answer |
|---|---|
| In one or two sentences, what do you understand by the term "scientist"? | (free text) |
| In one or two sentences, what do you understand by the term "physician-scientist"? | (free text) |
| In your opinion, what is the value of being a physician-scientist? | (free text) |
| Now that you have completed your miniproject, please rate your competency in the following areas.<br>Information on the meaning of each area is provided underneath. | |
| Area | Rating |
| Technical science skills[a]<br><br>[a.] Your ability to perform an experiment in the laboratory. For example, during your miniproject, you will learn how to perform confocal microscopy or cognitive testing. | Poor |
| | Fair |
| | Average |
| | Good |
| | Excellent |
| Critical assessment of science publications[b]<br><br>[b.] Your ability to read a scientific paper and interpret and critique its data, for example a paper that provides background for your miniproject. | Poor |
| | Fair |
| | Average |
| | Good |
| | Excellent |
| Please rate your competency in the following areas.<br>Information on the meaning of each area is provided underneath. | |
| Area | Rating |
| Time management[a]<br><br>[a.] Your ability to plan ahead to reach your targets, for example to complete a series of experiments in your miniproject. | Poor |
| | Fair |
| | Average |
| | Good |
| | Excellent |
| Organisational skills [b]<br><br>[b.] Your ability to schedule your tasks and your punctuality, for example, to use a particular piece of equipment that you need for your miniproject. | Poor |
| | Fair |
| | Average |
| | Good |
| | Excellent |
| Please rate your competency in the following areas.<br>Information on the meaning of each area is provided underneath. | |

| Area | Rating |
|---|---|
| Communication skills[a]<br><br>[a.] Your ability to communicate effectively and efficiently with colleagues, your superiors and the public in written or oral format, for example, your ability to troubleshoot with your superiors and your ability to use technical and/or simple language to explain your data. | Poor<br>Fair<br>Average<br>Good<br>Excellent |
| Presentation skills[b]<br><br>[b.] Your ability to present data to your colleagues, your superiors, to the public. For example, at the end of the program, all students will present their data in the multinational meeting, and all students will present their data and their experiences to the public during European Science night. | Poor<br>Fair<br>Average<br>Good<br>Excellent |
| Writing reports[c]<br><br>c. Your ability to write a report that is accurate, fair and based upon evidence, for example, a report on the data that you obtain during your miniproject. | Poor<br>Fair<br>Average<br>Good<br>Excellent |
| Please rate your competency in the following areas.<br>Information on the meaning of each area is provided underneath. | |
| Area | Rating |
| Ability to work in teams[a]<br><br>[a.] Your ability to reach agreement on tasks to do within a team and ability to complete that task, for example, during the public presentation during European Science night. | Poor<br>Fair<br>Average<br>Good<br>Excellent |
| Leadership skills[b]<br><br>[b.] Your willingness to take on responsibility and organise team-based tasks, for example, during the public presentation during European Science night. | Poor<br>Fair<br>Average<br>Good<br>Excellent |
| What supported your development of the competences described above? | (free text) |
| From your research experience, how much of a gain occurred in: | |
| Understanding how professionals work on real problems | No gain or very small gain<br>Small gain<br>Moderate gain<br>Large gain<br>Very large gain |
| Sense of contributing to a body of knowledge | No gain or very small gain<br>Small gain<br>Moderate gain<br>Large gain |

| | |
|---|---|
| | Very large gain |
| Understanding that scientific assertions require supporting evidence | No gain or very small gain |
| | Small gain |
| | Moderate gain |
| | Large gain |
| | Very large gain |
| Learning ethical conduct in your field | No gain or very small gain |
| | Small gain |
| | Moderate gain |
| | Large gain |
| | Very large gain |
| Self-confidence | No gain or very small gain |
| | Small gain |
| | Moderate gain |
| | Large gain |
| | Very large gain |
| In your opinion, what was the value of participating in this program? | (free text) |
| Evaluate your overall satisfaction with your miniproject experience. | I feel very dissatisfied |
| | I feel moderately dissatisfied |
| | I feel neutral about it |
| | I feel moderately satisfied |
| | I feel very satisfied |
| As a final question, how can we improve this research exchange program? | (free text) |

### Supplementary material 3

Based upon requirements identified by authors Monika Jürgenson, Andrea García-Llorca, Anu Sarv, Thor Eysteinsson and Miriam Hickey and upon published commentary (1).

**Post-1-year** follow-up questionnaire
*Mode: Confidential*

| Question | Answer |
|---|---|
| What is your ongoing science activity? | (free text) |

### Supplementary material 4

**Practical guidelines for organizing international research exchanges**

1) Advertising the program:

- Logo and communication:
    - Create a logo for the program, ensuring it's used in all official materials and communications (e.g., emails, posters, promotional materials).
- Introduction video:
    - Develop a short, engaging video to introduce the program's goals and benefits.
- Centralized website:
    - Build a comprehensive website that outlines research opportunities, program goals, eligibility, funding, application details, and daily/weekly lab commitments.
    - Partner universities should link to this central site.
- Early input:
    - Engage faculty administrators early for advice on managing exchange logistics and visiting student status.
    - Consult student council members for input on effective advertising tailored to the student population.
- Communication channels:
    - Consider face-to-face outreach and involving past participants to share their experiences.
- Virtual sessions:
    - Organize virtual sessions introducing faculty members from partner universities to give students more insight into the program.
- Application period:
    - Ensure adequate time for applications (1 month between initial invitation and deadline) and issue timely reminders.

2) Selection process:

- Conflict of interest:
    - Faculty members with a teaching relationship to applicants should not be involved in the selection process.
- Motivational letter:
    - Require students to submit a motivational letter as part of their application.
    - Establish clear rules regarding the use of AI in applications and use anti-plagiarism software if necessary.
- Second round of selection:
    - Consider a second round, where students present a science paper.
    - The paper should be selected and distributed by faculty, with clear guidelines on presentation and scoring.
    - Students must present their work independently, and faculty will score the presentations.
    - Consider allowing "back-up" participants in case of withdrawals.

3) Information packs:

- Content:

    - Provide selected students with a detailed information pack, including program specifics, pre-project training deadlines, accommodation options, local travel tips, and insurance information.
- Accommodation and travel:
    - Ensure, that if needed, faculty can help students with accommodation selection.
- Time commitment:
    - Reiterate the expected laboratory work hours and professionalism standards.
- ECT requirements:
    - Define conditions for awarding ECTs (if applicable).
- Input from past participants:
    - Involve past participants and local faculty in creating information relevant to day-to-day student experiences.

4) Funding:

- Funding clarity:
    - Clearly define what funding covers and the conditions for reimbursements.
- Sufficient but focused:
    - Ensure funding is sufficient but not designed to encourage students to treat the program as a vacation.
    - Align funding guidelines with schemes like ERASMUS.

5) Pre-exchange preparation:

- Dates and supervision:
    - Agree on the exchange dates with local faculty, ensuring proper on-site supervision.
    - Ensure all necessary experimental ethical approvals are in place prior to students' arrivals.
- Travel details:
    - Provide students with all necessary contact information (e.g., supervisor, emergency contacts) ahead of travel.
- Safety guidelines:
    - Offer comprehensive safety training before travel, including quizzes and signed safety agreements.
    - Also provide guidelines on clothing, laboratory footwear, and required protective equipment.
- Online learning platform:
    - Develop a course webpage with all program details and required safety and ethics training, accessible through the university's online platform (e.g., Moodle, Canvas, Blackboard).
- Communication skills:
    - Provide students with training on formal scientific communication and presenting to both scientific and lay audiences.

6) The exchange:

- Laboratory expectations:
    - Clearly define lab work expectations (e.g., hours, record-keeping).

    - Students should not start independent work until they demonstrate sufficient competence.
- Real-world projects:
    - Ensure projects are based on real-world research, with no guarantee of positive outcomes.
    - Students should be informed that projects may evolve as new results emerge.
- Project introduction:
    - Have students write a short introduction to their project within the first week.
- Reporting:
    - Require students to complete weekly progress reports, signed by faculty, to track activities and monitor professionalism.
    - A template progress report should be provided.
- Final report:
    - At the end of the project, students should submit a detailed project report that contextualizes their findings, critiques their work, and makes it accessible to peers and faculty for feedback.
- ECTs:
    - Consider awarding ECTs upon successful completion of the program and submission of required reports, with clear criteria for this recognition.

7) Assessing program effectiveness:

- Surveys:
    - Conduct surveys before, after, and a few months post-exchange to assess the impact of the program on students' science knowledge and their continued engagement in research.
- Dissemination:
    - Publish and share the findings of the surveys to measure the program's impact and improve future iterations.

8) Concluding the exchange:

- Debrief conference:
    - Organize a conference for all participating students and faculty to discuss their experiences, results, and program outcomes.
    - This can be virtual, hybrid, or in-person, depending on circumstances.

9) Open science involvement:

- Public engagement:
    - Involve students in open science events to share their research and experiences with the general public.
    - These events can foster community interest in scientific exchanges and the work students conducted during their program.

10) Program flexibility:

- Adapting to challenges:
    - Recognize that international exchange programs may face delays (e.g., due to COVID-19) and students should be flexible in adjusting to changes.

- Consider contingency plans and support for students who opt to wait for travel to become feasible.
